# Large-scale eQTL analyses in Atlantic salmon reveal persistent dosage compensation 100 million years after genome duplication

**DOI:** 10.64898/2025.12.15.694296

**Authors:** Célian Diblasi, Domniki Manousi, David Hazlerigg, Lars Grønvold, Nicola Jane Barson, Simen Rød Sandve, Marie Saitou

**Affiliations:** Department of Animal and Aquacultural Sciences, Section for Genome Biology, Faculty of Biosciences, Norwegian University of Life Sciences, 1433 Ås, Norway; The Roslin Institute and Royal (Dick) School of Veterinary Studies, University of Edinburgh Easter Bush, EH25 9RG, Midlothian, United Kingdom; Department of Arctic and Marine Biology, UiT The Arctic University of Norway, N-9037, Tromsø, Norway

## Abstract

Whole-genome duplication (WGD) through autopolyploidization has played a role in genome evolution across eukaryotes. A major consequence of WGD is the rewiring of gene regulatory networks, partly driven by selection on dosage balance. In multicellular organisms, evidence for dosage balance selection has relied on comparative patterns of duplicate gene retention and expression, with few studies directly examining regulatory architecture after WGD.

Here, we analysed a large-scale eQTL dataset from Atlantic salmon (Salmo salar), which experienced a WGD 100 million years ago. We found that trans-regulatory connections were enriched between duplicated regions, indicating long-term conservation of ancestral interchromosomal regulatory interactions. Overall, 230 duplicated genes (5%) shared eQTLs, suggesting conserved regulatory control. Moreover, 16 gene pairs showed compensatory expression effects mediated by a common regulator, consistent with predictions of the dosage balance hypothesis. These gene pairs were significantly enriched in recently rediploidized regions. Our results indicate long-term maintenance of dosage balance after WGD.

**Teaser:** Genetic regulation in Atlantic salmon shows that duplicated genes can remain dosage-balanced across 100 million years of evolution.

## Introduction

Whole genome duplication (WGD) is a major driving force in genome evolution across the eukaryote tree of life (*1–4*). Doubling of the genetic material results in functional redundancy and reduced selective constraints. This allows for mutations to accumulate (*1*, *5*) which impact trait evolution including gene regulatory phenotypes (*6–8*). However, our understanding of how WGDs impact the gene regulatory architecture through rewiring of cis- and trans-regulatory connections is only addressed in a few studies in yeast (*9*, *10*) and is still limited.

Gene expression levels evolve under selection for optimum ‘trait’ values (*11*, *12*). These optima change depending on (i) external factors such as the local environment and (ii) internal genomic factors such as imbalances in gene dosage that upsets the stoichiometric balance between RNA and/or protein levels (*13*, *14*). The genomic upheaval that follows WGDs, including rampant pseudogenization of redundant duplicate copies, must therefore spark selection to minimize stoichiometric imbalances. This could be achieved by selection to retain a balanced number of gene copies involved in dosage-sensitive interactions (*15*, *16*), selection on gene expression levels (*17*), or a combination. Studies of long-term evolutionary consequences of WGDs highlight that asymmetric evolution of ohnolog regulation is a common fate (*17*, *18*) resulting in one copy dominating expression. This is in line with a model whereby optimal expression levels are mostly resolved through selection on the sum of ohnolog expression levels, blind to which copy contributes to the expression trait value.

While numerous comparative transcriptomics studies have provided valuable insights into ohnolog regulatory evolution, our understanding of the genetic mechanisms in play is limited. To address this knowledge gap, associations between genetic variation and variation in gene expression levels, i.e. expression Quantitative Trait Loci (eQTLs), offers a rare opportunity to study patterns of conservation and divergence in gene regulatory mechanisms across duplicated genes (*19*, *20*). Here we leverage a large eQTL dataset from Atlantic salmon, a lineage which underwent autopolyploidization 100 Mya (*21*, *22*), to characterize divergence in gene regulatory network connections inferred by trans-eQTL associations. This lineage is particularly interesting due to its two waves of rediploidization, resulting in ancestral ohnolog resolution regions (AORe; ∼100mya) and lineage-specific ohnolog resolution regions (LORe; <50mya) (*21*, *22*).

## Results

### Trans-regulatory connections across homeologous regions are retained following WGD

To identify trans-regulatory associations, we used a recently published dataset of bulk-RNAseq from gill samples of ∼1000 Atlantic salmon raised on constant photoperiod, with matching genome wide SNP data for all individuals (*23*). Using this data, we identified 12,056 eQTL impacting the expression of 19,526 genes in gill tissue. Of these, 52% (5,951) were cis-eQTLs and 43% (5,022) were trans-eQTLs, with the remaining 5% (556) showing both types of interaction (**Figure 1A, B**). Note that the numbers of eQTL in this study differ from (*23*) due to the different sample sizes in the two analyses.

**Figure 1:**
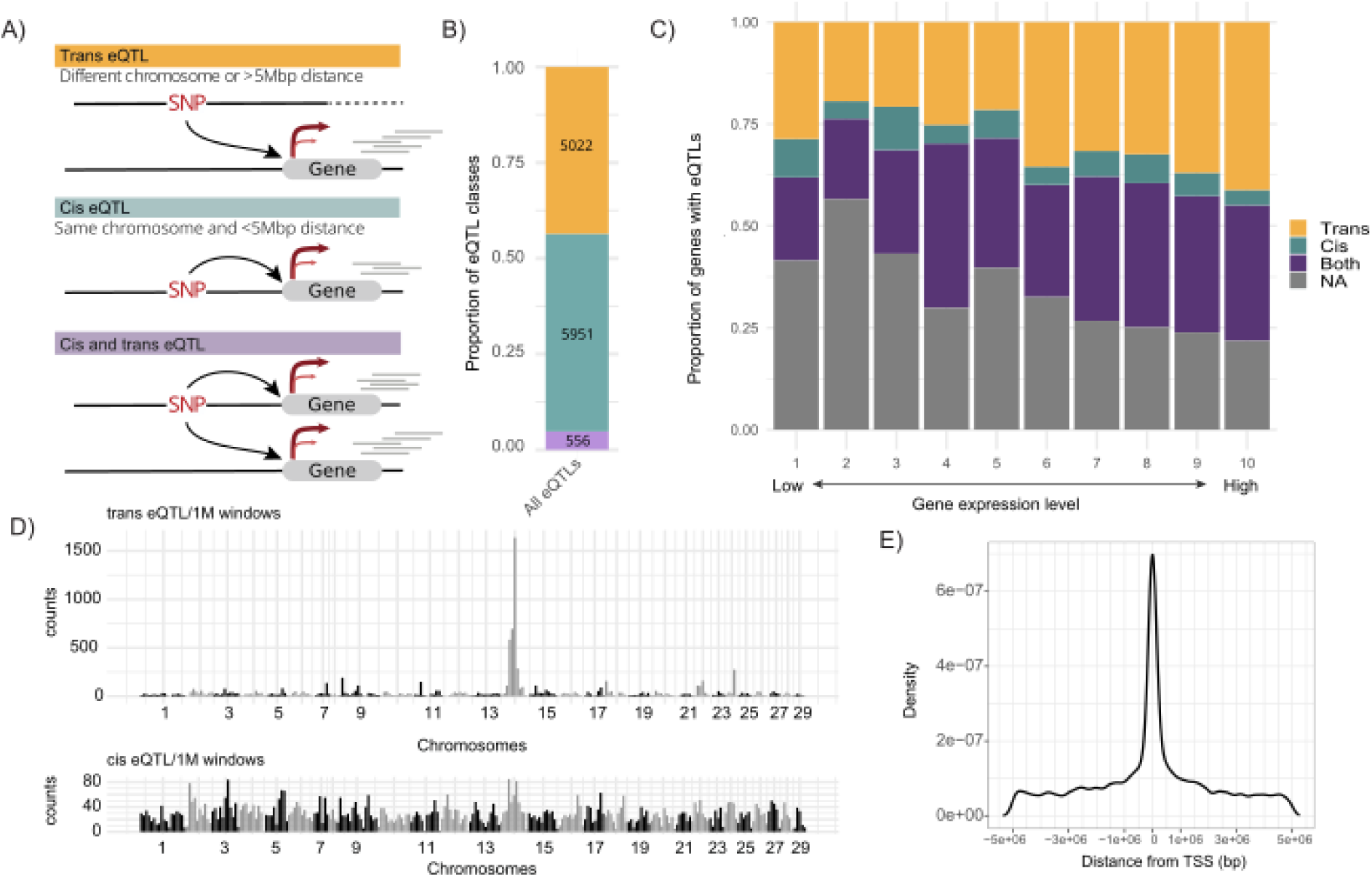
Overview of cis and trans eQTLs and their genomic distribution in Atlantic Salmon. A: Schematic illustration of different eQTL-gene relationships. **A:** Cis-eQTLs are defined as when SNP is located within 5 Mb of the gene it regulates. Trans-eQTLs are those where the SNP and gene are located more than 5 Mb apart or on different chromosomes. The third category involves eQTLs that show both cis and trans interactions. **B:** The proportion of identified eQTLs: 49% (5,887) are cis-eQTLs, 46% (5,549) are trans-eQTLs, and 5% (620) show both cis and trans interactions. **C:** Proportion of genes with cis, trans, both, or no eQTLs across 10 groups of genes, ranked by expression levels from low to high. Each group represents 10% of the genes, with group 1 having the lowest expression and group 10 the highest. **D:** Genome-wide distribution of trans-eQTLs (top) and cis-eQTLs (bottom) across chromosomes, divided into 1 Mb windows. Chromosome 14 shows a marked enrichment for trans-eQTLs (chi² = 29914, p-value < 2.2E-16), while no such enrichment is seen for cis-eQTLs. **E:** Density plot of distances between cis-eQTLs and the transcription start site (TSS) of the genes they regulate.

We estimated the average effect size to be 0.358 for cis-eQTLs and 0.379 for trans eQTL. Gene expression levels were correlated with the proportion of genes with an eQTL, as well as the proportion of trans-eQTLs (**Figure 1C**), most likely due to increased power to detect eQTLs. Highly expressed genes without detectable eQTLs were predominantly involved in ribosomal function (GO:0005840, FDR=2.8E-155), in line with the regulation of these genes being under strong purifying selection (*24–26*) (**Figure 1C**). Distribution of trans- and cis-eQTLs showed that chromosome 14 was strongly enriched for trans-eQTLs (chi² = 29914, p-value < 2.2E-16), while there was no such chromosomal bias present in cis-eQTLs associations (**Figure 1D**). Among the most significant cis-eQTLs for each gene, 35.3% were located within 500kb (10% of the cis-definition range) of the transcription start site (TSS) (**Figure 1E**).

### WGD and chromosomes fusions affect trans-eQTL connections

Immediately after a WGD occurs, all the gene regulatory connections are duplicated. However, during the subsequent rediploidization, regulatory connections may be lost or rewired independently among ohnolog copies (*16*, *27*). Trans connections typically refer to interactions between proteins, such as transcription factors, and genomic elements that are not necessarily immediately adjacent such as enhancers or distant regulatory sequences (*28*, *29*). It is thus reasonable to expect that when a genomic region is duplicated, these interacting elements may be able to bind to both copies of the sequence. Furthermore, following whole genome duplication (WGD), new trans connections can arise. These could in theory involve interactions between a duplicated gene and its ‘original’ cis-regulatory element landscape physically located on the duplicate copy chromosome. However, this is rather unlikely due to the inherent local nature and distance-dependent specificity of cis elements (*30*). New trans connections can also arise between genes and regulatory elements that become physically linked on the same chromosome as a result of chromosomal fusions. To investigate the trans connections landscape after WGD, we classified trans-eQTLs into the following four groups (**Figure 2A**): **(i)** Inter-chromosomal homeologs where the eQTL and target gene reside in homeologous regions (regions duplicated by WGD) on different chromosomes, **(ii)** inter-chromosomal non-homeologs where the eQTL and the target gene are located on different chromosomes but not in homeologous regions, **(iii)** intra-chromosome syntenic where eQTL and the target gene reside on the same chromosome within a syntenic block originating from the WGD >5 Mbp apart, and **(iv)** intra-chromosome non-syntenic where the eQTL and the target gene reside on the same chromosome as a result of chromosome fusion following WGD (*22*). A schematic representation of evolutionary events leading to these four different types of trans regulatory connections before and after WGD is shown in **Figure 2B**.

**Figure 2.**
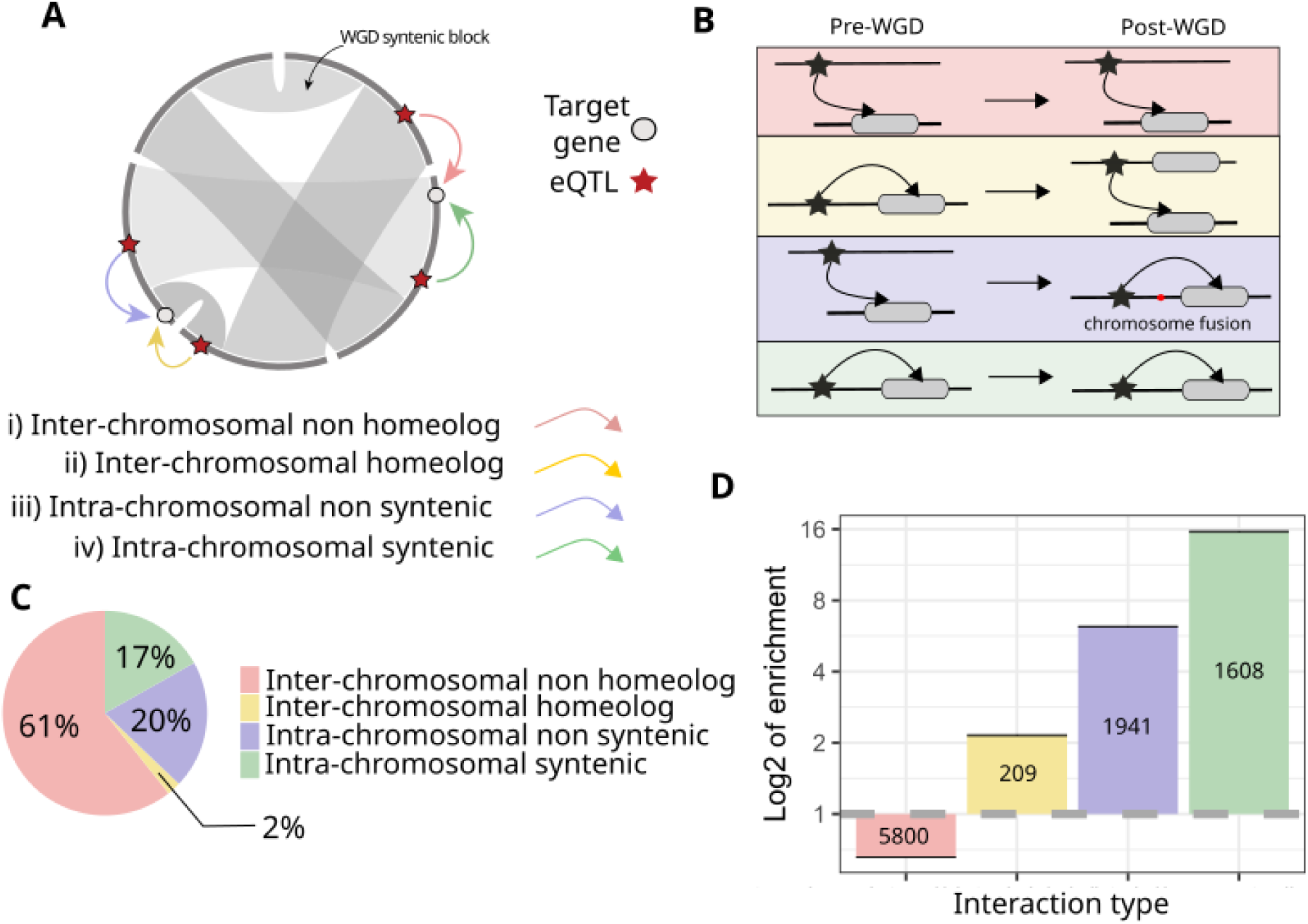
Types of trans-regulatory interactions in the context of WGD and their abundance. **A)** Four types of trans regulation: i: The eQTL SNP is associated with a gene on another chromosome that is not the homeologous region (inter-chromosome non-homeolog). ii: The eQTL SNP is associated with a gene on the homeologous region on a duplicate chromosome originating from WGD (inter-chromosome homeolog). iii: The eQTL SNP is associated with a gene on the same chromosome outside the syntenic region (intra-chromosome non-syntenic) iv: The eQTL SNP is associated with a gene in the same syntenic region (intra-chromosome syntenic). **B)**. Schematic representation of evolutionary events leading to different types of trans regulation (Left: before WGD, right: after WGD). **C)**. Proportion of the different types of trans regulation in observed connections. **D)**. Enrichment or depletion of each type of trans-regulatory interaction compared to randomly shuffled interactions. Enrichment values above 1 indicate that the interaction type is more frequent than expected by chance (**see Methods**), with the number of observed connections indicated within the bars. Standard deviation is shown in black.

Among the four trans-regulatory categories, most (61%) were inter-chromosomal non-homeolog, while 20% were intra-chromosomal non-syntenic, 17% were intra-chromosomal non-syntenic, and only 2% were inter-chromosomal homeologs (**Figure 2C**). Compared to randomized data (**see Materials and Methods**) this represents a 13.7-fold enrichment of intra-chromosomal syntenic trans-eQTL connections (category iii) and a 6-fold enrichment of intra-chromosomal non-syntenic trans-eQTL connections (category iv) (**Figure 2D**). These results suggest that trans-regulatory connections within the same chromosomes tend to be retained, and that new regulatory connections are being established between variants and genes within newly fused chromosomes. We observed that the distance between eQTLs and the target genes were higher for intra-chromosomal non-syntenic trans-eQTL connections (category iv) than for intra-chromosomal syntenic connections (category iii), even after correcting for differences in the maximum eQTL-gene distance between the two categories. This result confirmed that the enrichment of intra-chromosomal non-syntenic trans connections is not a technical artifact arising from the distance cutoff (**Figure S1**).

With respect to inter-chromosomal relationships, we observed two times more inter-chromosomal homeologous eQTLs target gene connections (category i) than expected by chance (**Figure 2D**), which suggest that regulatory elements, originally located on the same chromosome as their associated genes before WGD, continue to influence the expression of their original target genes after being separated into different chromosomes by WGD (**Figure 2 A, B**). It has previously been shown that transcription factor genes are preferentially retained after WGD (vertebrates: Kassahn et al. 2009, paramecia : McGrath et al. 2014, plants: Blanc & Wolfe 2004), however, we did not observe significant enrichment for any GO term, including transcription factors, for genes in inter-chromosomal homeologous trans-eQTL connections (FDR cutoff = 0.05).

To test the impact of rediploidization timing on trans-eQTL evolution we next compared early (LORe, <50 Mya) vs late (AORe, ∼100 Mya) rediploidizing regions. Inter-chromosomal homeologous trans connections (category ii) were more frequently retained in LORe compared to AORe regions (chi² = 95.6, p-value = 2.2e-16) (**Figure S2**).

### Shared trans-eQTLs are associated with higher gene expression correlation

Immediately following a WGD, newly duplicated ohnologs share identical “ancestral” regulatory networks. This includes the same regulatory molecules (both with indirect and direct effects) and identical cis-regulatory motifs where the direct-acting regulatory molecules can interact with promoters and enhancers. Through time, these duplicated regulatory networks will diverge, e.g. through divergence in the cis-regulatory landscape, driving gene regulatory divergence between ohnologs.

To assess the level of ohnolog regulatory divergence, we classified ohnolog pairs into three different ‘ohnolog eQTL divergence classes’ **(Figure 3A)**; **(i)** ohnologs sharing an eQTL, **(ii)** ohnologs with distinct eQTLs, **(iii)** ohnologs where only one copy has an eQTL, and **(iv)** ohnologs without eQTLs. To reduce the risk of misclassifying ohnolog pairs due to statistical noise in lead SNP selection, we applied fine-mapping with functionally informed priors (**see Methods, Figure S3**). This method improves the resolution of causal variant identification by estimating the likelihood that a variant directly affects gene regulation rather than merely reflecting statistical association. For each gene, we defined causal eQTLs as fine-mapped cis or trans variants with non-zero maximum posterior inclusion probability (PIP) (**Figure S4**).

**Figure 3:**
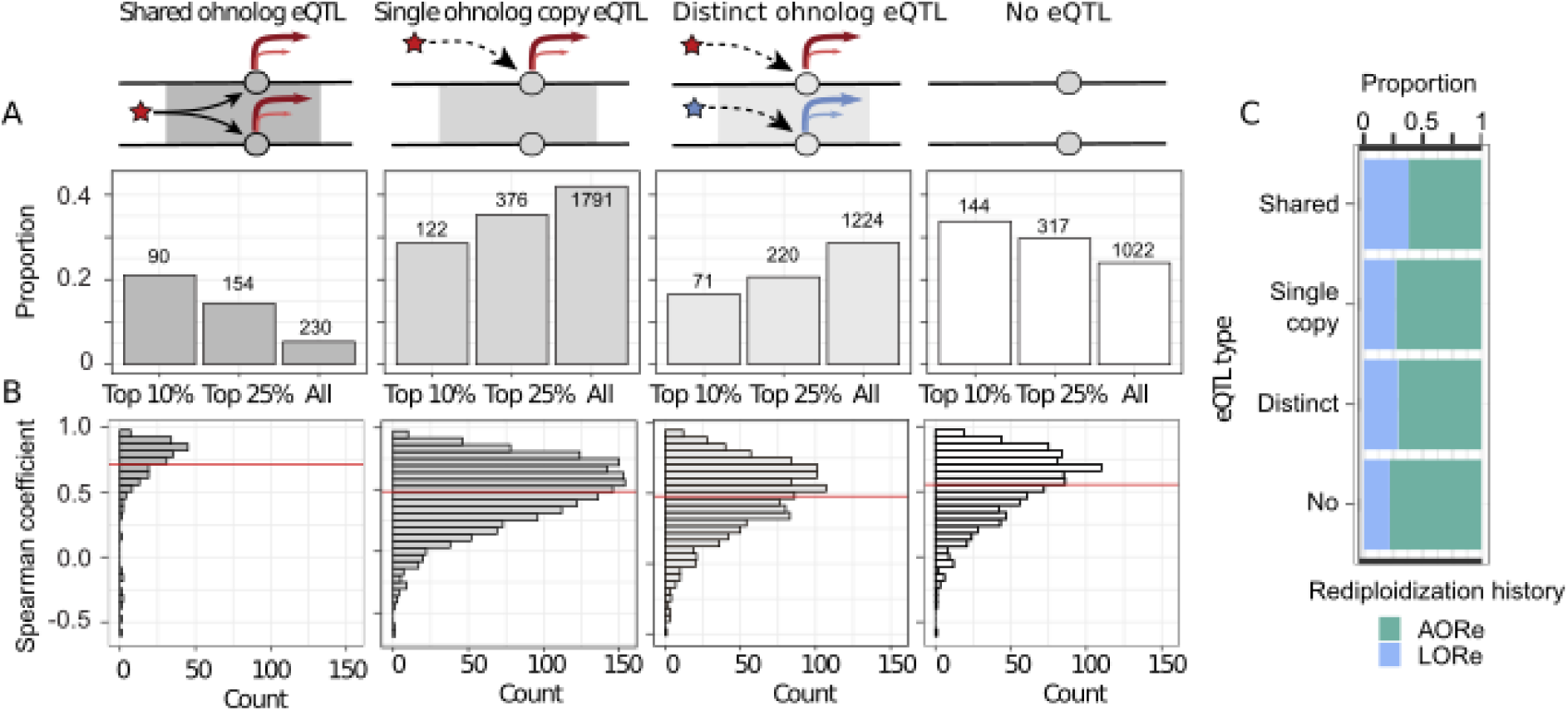
Ohnolog regulation patterns and expression correlations between ohnolog gene pairs. **A.** Four regulation states are defined: **i:** Both copies of a gene are regulated by the same eQTL (shared eQTL). **ii:** One of the two copies is regulated by an eQTL (Single copy eQTL. **iii**: Both copies are regulated by different eQTLs (distinct eQTL). **iv:** None of the copies are regulated by an eQTL (no eQTL). The proportion of ohnolog pairs in each regulatory category is shown for all pairs, as well as for the top 10% and 25% of pairs with the highest correlation in gene expression. **B:** Spearman correlation coefficients of gene expression between ohnolog pairs for each regulatory category. The horizontal red line represents the mean correlation. **C:** The proportion of pairs in the AORe (green) and LORe (blue) region depends on their regulatory context.

Our analysis of the fine-mapped eQTLs revealed that 5% of the ohnolog gene pairs (230/4,267) share at least one fine-mapped eQTL. For those genes, we identified a GO enrichment in functions related to mitochondrial pathways and respiration (**Figure S5**). Among these 230 ohnolog gene pairs with shared eQTL, 229 pairs were regulated by trans-eQTLs, with only one pair sharing both cis- and trans-eQTLs. Conversely, 1,224 ohnolog pairs (29%) had distinct eQTLs for each gene copy (**Figure 3B**). If shared eQTLs actually reflect that ohnologs have more conserved (and similar) regulatory networks, one expectation is that these ohnologs should have higher gene expression correlation compared to those that do not share eQTLs. To test this, we used the absolute Spearman’s correlation coefficient for each ohnolog pair across all samples, to only account for correlation strength. In line with this expectation, ohnolog pairs with shared eQTLs exhibited significantly higher correlation (Spearman’s rho: 0.74 ± 0.01) compared to pairs with distinct eQTLs (rho: 0.49 ± 0.01), single-copy eQTLs (rho: 0.50 ± 0.01), or no eQTLs (rho: 0.56 ± 0.01) (**Figure 3B**). Moreover, the distribution of expression correlation for shared eQTL pairs was significantly (p < 2E-16) different from the other regulation categories (**Table S1**). These trends were still present when considering only genes within LORe (rho: 0.70; 0.52; 0,53; 0.56, respectively) or AORe (rho: 0.75; 0.49; 0,48; 0.56, respectively) regions. In addition, we found that the mean absolute correlation of observed ohnolog gene pairs (rho: 0.52 ± 0.004) was much higher than for randomly shuffled gene pairs (rho: 0.30 ± 0.003), confirming the strong positive correlation between ohnolog gene pairs.

Because divergence time is associated with regulatory network divergence of gene duplicates (*19*, *34*) we asked if the proportion of shared trans-eQTLs was associated with the timing of rediploidization (*22*). Indeed, more ohnolog regulatory divergence (i.e. distinct and single classes) was observed in ohnologs residing in early rediplodized regions (AORe) compared to late rediploidized regions (LORe) (chi² = 9.7, p-value = 0.02) (**Figure 3C**).

Taken together we find strong evidence for long term conservation of regulatory networks among ohnologs. However, the majority of ohnologs do not share eQTLs, consistent with previous findings of asymmetric ohnolog expression evolution dominating after the salmonid WGD (*17*).

### Genotype-dependent dosage compensation in ohnolog gene pairs

The vast majority of ohnolog pairs with shared eQTLs had genetic variants impacting gene expression in the same direction in both ohnolog copies. However, for a small proportion of ohnologs (7 ohnologs gene pairs) the alleles of the shared eQTL had opposite effects on the gene expression (**Figure S6**), indicating dynamic dosage compensation mechanisms (**Figure 4A**).

**Figure 4:**
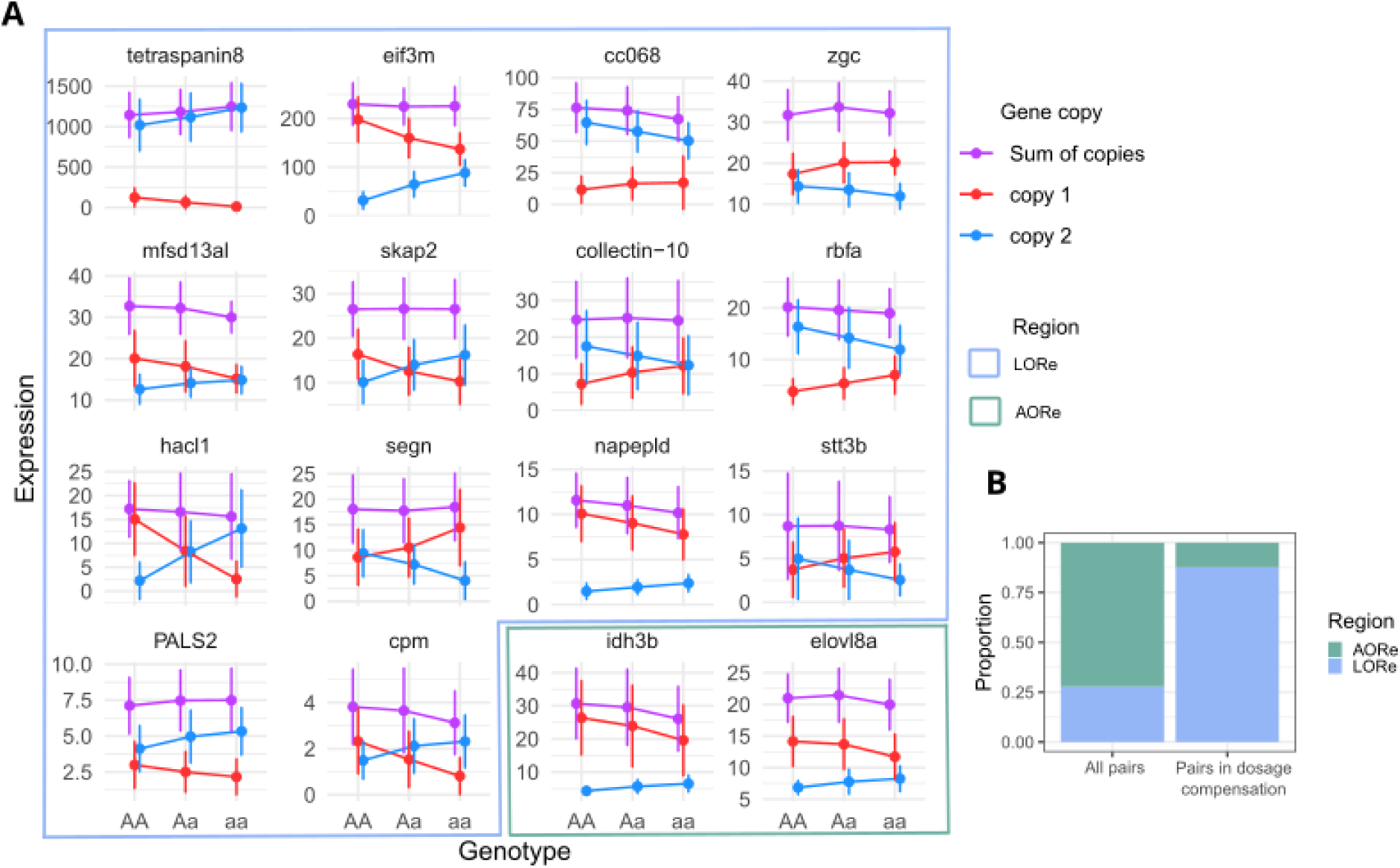
Gene expression levels of dosage compensated ohnolog according to their eQTL genotype. **A:** Mean expression levels (in TPM) for the 16 ohnolog pairs with shared eQTLs and negative expression correlations, with their standard deviation. The expression is grouped by the genotype of the shared eQTL, which are represented by aa, Aa, and AA. Red lines represent Copy 1, blue lines represent Copy 2, and purple lines represent the sum of both copies’ expressions. **B:** Proportion of dosage compensated ohnolog (among the 16 found) in AORe and LORe region versus all gene pairs.

To investigate this phenomenon in more detail we reanalysed the data using less stringent criteria for identifying shared eQTLs across ohnolog pairs showing negative expression correlation. Specifically, some shared eQTLs could have been misclassified as distinct eQTLs due to the selection of different eQTLs as fine mapped eQTL for each gene copy, despite the eQTLs being in linkage disequilibrium and tagging a shared, true causal variant. Out of 53 negatively correlated ohnolog pairs that were previously defined as having distinct trans-eQTLs, 9 pairs were redefined as having shared eQTLs (**Figure S3**), increasing the total number of negatively correlated gene pairs with shared eQTLs to 16 (0.4% of the 4,267 pairs). These are referred to as genotype-dependent dosage compensated ohnologs (**Figure 4A**).

For the majority (11/16 = 69%) of the dosage compensated ohnologs the expression sum of both copies did not differ across eQTL genotypes, while for the others the expression varied from 5% to 19% of the total expression range (**Table S2**).These genes include several highly conserved housekeeping genes such as Eukaryotic Translation Initiation Factor 3 Subunit M (*eif3m*), ribosome binding factor A (*rbfa*), and N-Acyl Phosphatidylethanolamine Phospholipase D (*napepld*). In most genotype-dependent dosage compensated ohnologs (9/16 = 56%), one copy consistently showed much lower expression compared to the other (e.g *tetraspanin8*), while in four ohnolog pairs this was not the case (e.g. *skap2* in **Figure 4A**) (**Table S2**). Additionally, we observed a significant enrichment of dosage compensated ohnolog in LORe regions compared to AORe regions (chi²=25, p-value=4.7E-7, 3.16-fold enrichment) **(Figure 4B).**

## Discussion

### Conserved trans-regulation across 100 million years

Here, we studied the evolution of gene regulation following WGD using a large eQTL dataset from Atlantic salmon gill. We found that most ohnologs have distinct eQTLs, in line with other studies demonstrating largely asymmetric gene expression following WGD in vertebrates (*17*, *18*). However, we found 229 ohnolog pairs with conservation of trans-regulatory connections, across 100-50 million years. The ohnologs that sharing trans-eQTLs exhibited the highest expression correlation across individual gill samples (**Figure 3**), as has been shown in recently duplicated genes in allopolyploid plants (*35*). This elevated expression correlation also indicates a higher level of gene-regulatory network conservation compared to other ohnologs. This pattern cannot be exclusively explained by ohnolog divergence time (i.e. rediploidization history), as the difference persists within the divergence time category. Hence, we argue that the shared eQTLs and strong co-expression of these 229 ohnolog gene pairs reflects strong purifying selection on gene regulatory networks. The presence of an important proportion of genes involved in respiration and energy pathway, which are known to be under stabilizing selection (*17*, *36*), further support this hypothesis.

### Selection on gene dosage constraints is heavily biased towards LORe regions

A small subset of ohnolog pairs (16 out of 4,267), most of which encode housekeeping genes, shared trans-eQTLs but exhibited negatively correlated expression levels (**Figure 4**). In addition, the expression levels summed across ohnologs and genotypes were often identical or varied only little (**Figure 4**). We interpret this pattern as evidence for genotype-dependent dosage compensation, a pattern expected when duplicated genes are at least partly functionally redundant and there is stabilizing selection on their combined expression level.

Two gene-regulatory mechanisms could in theory give rise to genotype-dependent dosage compensation. One possibility is feedback regulation, in which excessive transcript levels from a gene directly or indirectly repress its own expression. In duplicated genes, such feedback could act asymmetrically, with one copy being more sensitive to repression, and the strength or direction of this effect varying with genotype (*37*, *38*). Alternatively, ohnologs could have evolved opposite regulatory outputs from the same trans-acting factor. In this scenario, a transcription factor might function as a repressor and activator, depending on the cis-regulatory element context. This mechanism is in line with recent work in yeast showing that many transcription factors act as both repressors and activators depending on promoter context (*39*). Although this seems like a straightforward explanation, the high sequence similarity between duplicates makes it uncertain to occur.

Notably, almost all ohnolog pairs with genotype-dependent dosage compensation signatures were located in late rediploidizing (LORe) regions, where sequence divergence is minimal and the encoded proteins are likely to be identical or nearly identical (*21*, *40*). The phenomenon of genotype-dependent dosage compensation itself is expected to arise when purifying selection acts on the sum of ohnolog gene expression and ohnologs are functionally redundant. However, the enrichment of this phenomenon in LORe is likely to be a consequence of the low “divergence time” of genes in those regions, thus having less chances to accumulate mutations that decouples their regulatory network interactions.

### Impact of genome structure evolution on gene regulatory connections after WGD

During rediploidization following WGD, gene regulatory connections are gradually lost or rewired (*41*, *42*). Here, we examined this process in the context of post-WGD karyotype evolution. Consistent with previous studies, we observed a strong enrichment of gene regulatory connections within chromosomes (*43*, *44*), potentially reflecting the facilitation of regulatory interactions by 3D chromatin conformation (*45*). This enrichment was higher (∼16-fold) in chromosomal regions with no karyotype reorganization since the WGD compared to those that have undergone more recent chromosome fusions (∼7-fold). One possible explanation for this phenomenon is that post-WGD chromosome fusions disrupted ancestral long-range cis-regulatory interactions between enhancers and promoters. Alternatively, fusions could also interrupt trans-regulatory interactions mediated by transcription factor activity. In humans, interacting proteins, complexes, and pathways tend to be non-randomly co-located within chromosomes (*46*, *47*), a pattern thought to arise from selection maintaining genes with shared functional properties in close physical proximity, and thereby maintain similar regulatory inputs from the surrounding enhancer landscapes. The lower enrichment of regulatory connections between recently fused chromosomal regions may therefore reflect the disruption of ancient hubs of co localized functionally related genes.

An additional pattern of interest is the enrichment of inter-chromosomal homeologous interactions. Although these connections constitute only about 2% of all observed trans-regulatory interactions (209 connections), they occur approximately two-fold more often than expected by chance. A possible explanation is that they represent physical contacts between duplicated chromosomes, potentially maintained by higher-order nuclear architecture since the WGD. Such contacts could preserve ancestral regulatory relationships even after the two homeologous regions were separated onto different chromosomes during rediploidization. The observed enrichment of these connections in LORe regions is consistent with reduced sequence divergence allowing the retention of ancestral cis-regulatory compatibility. These interactions would then reflect ancient cis-regulatory elements that, after WGD, became positioned on different chromosomes yet continue to exert trans effects on their original target genes. Further investigation of 3D genome conformation data (e.g., Hi-C) to directly assess spatial proximity between homeologous chromosome pairs would be needed to confirm this hypothesis.

Our study provides insights into how WGD shapes gene regulatory evolution, linking genomic duplication events to long-term regulatory dynamics. We identified sixteen dosage-compensated genes under purifying selection, indicating that balanced expression levels have been maintained by evolutionary constraints for over 100 million years. We also found that newly formed regulatory connections are enriched in intra-chromosomal interactions, suggesting an important contribution of 3D chromatin conformation to the emergence of rewired regulatory links after WGD.

Building on these insights into regulatory rewiring, further studies could provide a more comprehensive understanding of these phenomena. The present study relied on RNAseq data from a single tissue, and because gene expression patterns can vary across tissues, incorporating additional tissue types in future work would help determine the degree of tissue or cell-type specificity or generality in these regulatory responses. Moreover, experimental validation of dosage compensation, for example through gene editing, would offer deeper mechanistic insight. This was not feasible in the present study due to the extremely high sequence similarity between LORe-embedded duplicates. The uses of different Atlantic salmon lineages having accumulated sequence divergence could eventually allow the targeting of duplicates for an experimental setup.

Taken together, our results contribute to the understanding of how WGD influences the evolution of trans-regulatory connections and the long-term maintenance of gene dosage compensation.

## Material and methods

### Ethical statement

The experiment adhered to EU regulations for the protection of animals used in research (Directive 2010/63/EU). All necessary precautions were taken to reduce pain and discomfort. The Norwegian Food and Safety Authority approved the experiment (FOTS ID: 25658).

### Fish samples

For a detailed explanation of the fish rearing process, refer to Gjerde et al. (2024). Briefly, three groups of approximately 3000 juvenile Atlantic salmon (*Salmo salar*) were subjected to different photoperiods: LL (continuous 24-hour light), 12:12 (12 hours light / 12 hours dark), and 8:16 (8 hours light / 16 hours dark). The LL group was maintained under continuous light from hatching until transfer to seawater (SW). The 8:16 group was reared under continuous light until they reached a body mass of approximately 50g, after which they were exposed to a short photoperiod (8L:16D) for 6 weeks, followed by 8 more weeks of continuous light before SW transfer at a body mass of around 100g. The 12:12 group experienced 12 hours of daylight and 12 hours of darkness for 6 weeks, followed by 8 weeks of continuous light prior to SW transfer. Gill tissues were collected one week before seawater transfer.

For this study, we focused on the 906 individuals reared under the LL condition. To perform imputation for these fish, we used genotyping data from a 66K SNP array and whole genome sequencing (WGS) data from their 112 parent fish.

### Whole genome sequencing

Initially, we identified 9.4 million SNP variants through whole genome resequencing of 112 parent fish using the GATK pipeline gatk4-spark:4.3.0.0 (*49*). After quality filtering with PLINK v1.9 (*50*) based on Hardy-Weinberg Equilibrium (HWE p-value < 1E-8), minor allele frequency (MAF < 0.01), and genotype missingness thresholds per individual and SNP variant (10% and 5%, respectively), we retained 108 individuals and 5.8 million SNPs for further analysis.

### Genotyping array

Fish were genotyped using a 66K SNP Affymetrix array [Salmow01]. Physical coordinates of array markers were re-assigned according to the newer Atlantic salmon genome reference (GenBank accession: GCA_905237065.2) and variants were quality-controlled using PLINK v1.9 (*50*, *51*). Quality filtering for genotypes used the following thresholds: HWE p-value < 1E-6, MAF < 0.01. Samples with more than 10% of their genotypes missing were removed, as well as variants missing in more than 3% of the total sample. A total of 2970 samples and 48,563 SNP were retained after quality filtering.

### Genotype imputation

To improve the SNP coverage for the 906 LL condition fish, we performed genotype imputation using the Beagle 5.2 software (*52*, *53*) leveraging the WGS data from the 112 parent fish. In order to identify SNP with high inference confidence, a five-fold cross-validation method was applied, by dividing the parent fish into five groups. The genotypes missing from the SNP array were masked in 20% of the individuals and imputed using the remaining 80%. Imputation accuracy was assessed as the square of Pearson’s correlation (R²) between the true and imputed genotypes, averaged across the five validation results. This process resulted in the retention of 412,069 SNPs with an R² > 0.80. After further quality filtering of imputed SNP based on HWE deviations (HWE p-value < 1E-8) using PLINK v1.9, the final imputed dataset consisted of 366,437 SNP markers, in addition to the original 48,563 SNPs from the genotyping array.

### RNA-sequencing

For detailed methods of RNA sequencing, see Grønvold et al. (2024). Briefly, gill biopsies were flash frozen, and RNA was extracted using the RNeasy Fibrous Tissue Mini kit (QIAGEN). Libraries were prepared and sequenced with 2x150 bp paired-end Illumina sequencing.

Before transcript quantification, adaptor sequences were removed using fastp version 0.23.2 (*55*). Read mapping and transcript quantification were conducted with Salmon version 1.1.0 (*56*), utilizing the Atlantic salmon transcriptome annotation from the Ssal_v3.1 genome assembly (Ssal_v3.1, GCA_905237065.2) from Ensembl. To prevent misalignment of reads to similar but unannotated regions, a transcriptome index file was used as a decoy sequence, applying the selective mapping mode (*57*). The --keepDuplicates and --gcBias options were used to address high sequence similarity between duplicated genes resulting from the salmonid genome duplication (*21*) and to correct for fragment-level GC biases, respectively. Gene-level mRNA expression was calculated by summing raw read counts and normalizing them into transcripts per million (TPM) using the R package tximport (*58*).

### eQTL analysis

We combined genotype and gene expression datasets using QTLTools (version 06 May 2020) (*59*) to identify eQTLs. Covariates including Father, Mother, Sex, Tank, Weight, Body length, Conditional factor, Smolt status, and ten genomic principal components (PCs) were regressed out to control for genetic relatedness and batch effects. This analysis resulted in the identification of cis-eQTLs (SNP-gene distance ≤ 5 Mb, FDR < 0.05) and trans-eQTLs (SNP-gene distance > 5 Mb or located on different chromosomes, FDR < 0.05) (**Figure 1**).

### Categorization of eQTL- gene connections in the context of WGD

WGD-derived synteny regions in Atlantic salmon genome were retrieved from https://salmobase.org/. Each trans-eQTL-gene pair was categorized into one of four groups based on the location of the eQTL and the gene, as detailed in the results section (**Figure 2**). To assess the statistical significance of the observed trans-eQTL connections, we generated a random expectation by shuffling the eQTLs and their associated genes for all identified trans-eQTL pairs. This randomization process was repeated 1,000 times.

To further analyse the differences in retention of inter-chromosomal homeologous connections between AORe and LORe regions, we first calculated the observed connection density by dividing the number of connections by the number of base pairs (bp) covered by each region. We then computed the expected density through 1,000 iterations of random shuffling for both AORe and LORe regions. Finally, a chi-square test was conducted to compare the observed and expected densities of connections between these regions.

### Fine-mapping with functional annotation

In the previous eQTL analysis on individual genes (**Figure 2**), we used the lead SNP for each gene. Next, we aimed to determine whether each ohnolog gene pair has a common eQTL, or distinct eQTLs (**Figure 3**). Using the lead SNPs naively could incorrectly suggest that a gene pair is associated with different eQTLs simply due to statistical noise, while they actually share a common genetic basis.

To address potential misclassification, we conducted fine-mapping using SparsePro (*60*) to identify putatively causal variants for cis and trans eQTLs separately. p-values from the QTLtools output, representing the association between genotypes and gene expression, were converted to Z-scores and used as input for SparsePro. For genes with either cis or trans eQTL SNPs, we allowed up to 10 credible sets (L = 10) in the fine-mapping process.

We applied fine-mapping with functionally informed priors, incorporating exonic regions and non-coding RNA locations from Ensembl Ssal_v3 annotation, and transcription factors retrieved from orthologous transcription factors from *Esox lucius*. For each gene, we defined its causal eQTL as the fine-mapped cis or trans variant with a non-zero maximum posterior inclusion probability (PIP), following an exploration of different PIP cutoffs for eQTL definition (**Figure S3 and S4**).

### Ohnologs expression analysis

Information from ohnologs genes were obtained using synteny information available at https://gitlab.com/sandve-lab/defining_duplicates/. Some gene pairs (ENSSSAG00000045167/ ENSSSAG00000001778 ; ENSSSAG00000044129/ ENSSSAG00000117896 ; ENSSSAG00000090152/ ENSSSAG00000110471) with no AORe/LORe information were manually assigned to LORe/AORe region using WGD-derived synteny region information. All ohnologs were classified among 4 categories depending on their regulator, as explained in the results section (**Figure 3**). For each gene duplicate, if the credible sets of two genes have at least one overlapping fine-mapped eQTL with non-zero posterior probabilities, we define the overlapping eQTL as the “shared” causal eQTL. Otherwise, they were classified into the three other categories, depending on if the two copies were regulated by two distinct eQTLs, or only one copy was regulated, or neither copy was regulated. Gene expression data was then used to compute spearman coefficients. We compared the rho of each distribution using absolute Spearman correlation values in order to investigate the strength of the association in expression of the pairs, regardless of its direction. Since not all gene pairs have recorded expression, these analyses were performed on 4267 pairs.

### Revisiting the classification of eQTLs for gene pairs with negative expression correlation

We initially identified nine ohnolog gene pairs that shared eQTLs and exhibited negatively correlated expressions, where higher expression of one copy was linked to lower expression of the other. However, upon closer examination of Ensembl and NCBI gene annotations, we discovered some discrepancies in the two databases: in some cases, a single gene in Ensembl corresponded to two distinct genes in NCBI (**Figure S7**). To resolve this, we reassessed the expression correlation using NCBI annotations and found that these ohnologs pairs actually showed positive correlations. Consequently, we excluded these pairs from the set of negatively correlated ohnologs.

This finding prompted us to further investigate the mechanisms of regulatory divergence following whole genome duplication (WGD), seeking to capture additional gene pairs that might have been overlooked. We also hypothesized that our fine-mapping procedure may have been overly stringent in defining shared eQTLs. Specifically, gene pairs might have been misclassified as having distinct eQTLs if the selected credible sets differed, especially when eQTLs were in linkage disequilibrium and tagging a common causal variant. To address this, we revisited the QTLtools results for 53 negatively correlated ohnolog pairs initially categorized as having distinct eQTLs. We reviewed all significant gene-eQTL associations (nominal p < 0.05) prior to fine-mapping (*59*, *60*). If the fine-mapped eQTL in one gene copy was also a significant eQTL for the other gene copy, we reclassified that pair as having a shared eQTL for the negative correlation analysis (**Figure S3**). This reanalysis recaptured 9 ohnolog pairs from the “distinct eQTL” category, bringing the total number of shared eQTL pairs with negative correlations to 16.

### Gene ontology enrichment

Gene Ontology (GO) enrichment analyses were performed using ShinyGO (V 0.80, http://bioinformatics.sdstate.edu/go/). For each gene set of interest, enrichment was tested using the default hypergeometric framework with Benjamini–Hochberg correction for multiple testing. Terms with an FDR < 0.05 were considered significantly enriched. The background gene universe was defined all registered genes in the default setting, unless specified otherwise. GO enrichment analyses were performed for specific gene sets, including:

i. highly expressed genes without detectable eQTLs,
ii. gene overlapped by Inter-chromosomal-homeologous trans eQTL,
iii. ohnolog pairs sharing at least one fine-mapped eQTL. (Gene set: the 4267 ohnologs pairs, singletons being ignored).

## Acknowledgements

We acknowledge the use of the Orion computing cluster at the Norwegian University of Life Sciences (NMBU). We thank Torgeir Rhoden Hvidsten from NMBU-KBM for his advice on expression network analysis. We thank Jun Soung Kwak from Kongju National University for his help on investigating the possibility to design and experiment to test the dosage compensation effect.

## Author contribution

The study was conceived by SRS, DH, CD, MS. Data was generated by DH and SRS. Bioinformatic and computational analyses were conducted by CD, DM, GG, LG and MS. Interpretation of the results were done by all authors. Drafting paper was done by CD, MS, LG, SRS, NJB. Manuscript editing was done by all authors.

## Data and Code availability

The Illumina sequencing data from RNAseq and whole genome resequencing are available in the European Nucleotide Archive under the project PRJEB47441. The code and output from bioinformatics pipelines used to generate all the figures and downstream analyses of data in R is available at https://github.com/celiand/eQTL_WGD_salmo.

