## Supplementary materials for "Large-scale eQTL analyses in Atlantic salmon reveal persistent dosage compensation 100 million years after genome duplication"

Célian Diblasi *et al*.

**This PDF file includes:**

Figs. S1 to S7

Tables S1 to S2

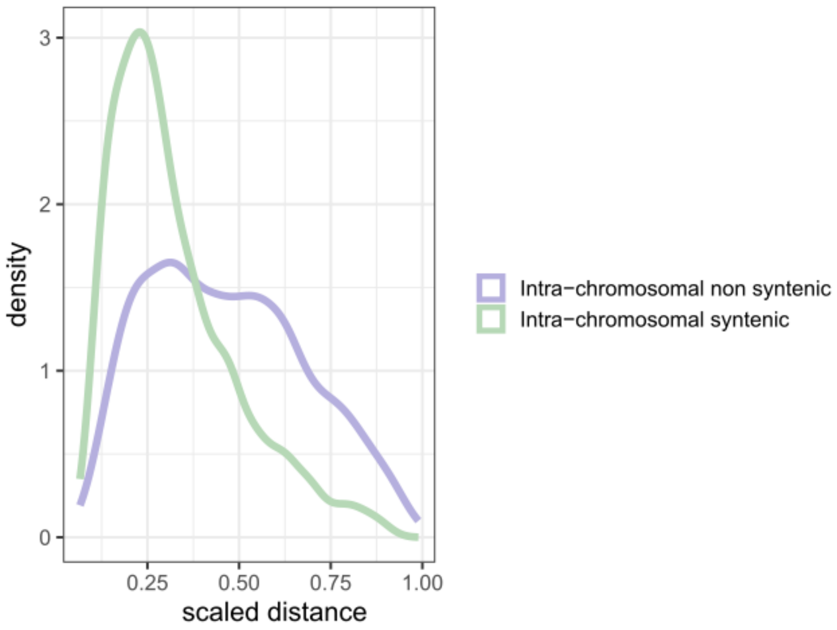


**Figure S1: Scaled physical distance between the trans eQTL and the regulated gene**

The distance between the eQTL and the gene is scaled by it, dividing by the maximum range possible for the eQTL-gene connection in the region where it's located. Since intra chromosome non syntenic connections can cover a greater range (mean=47 024 214 bp, max=161 086 148 bp) than intra chromosome syntenic connections (mean=34 021 492 bp, max=56 025 461 bp), an identical raw distance will generally be a higher scaled distance in intra chromosome syntenic connections than in intra chromosome non syntenic connections, which could bias our results. However, even with this potential bias, intra chromosome non syntenic connections are larger with scaling (mean=0.47) than intra chromosome syntenic connections (mean=0.28).

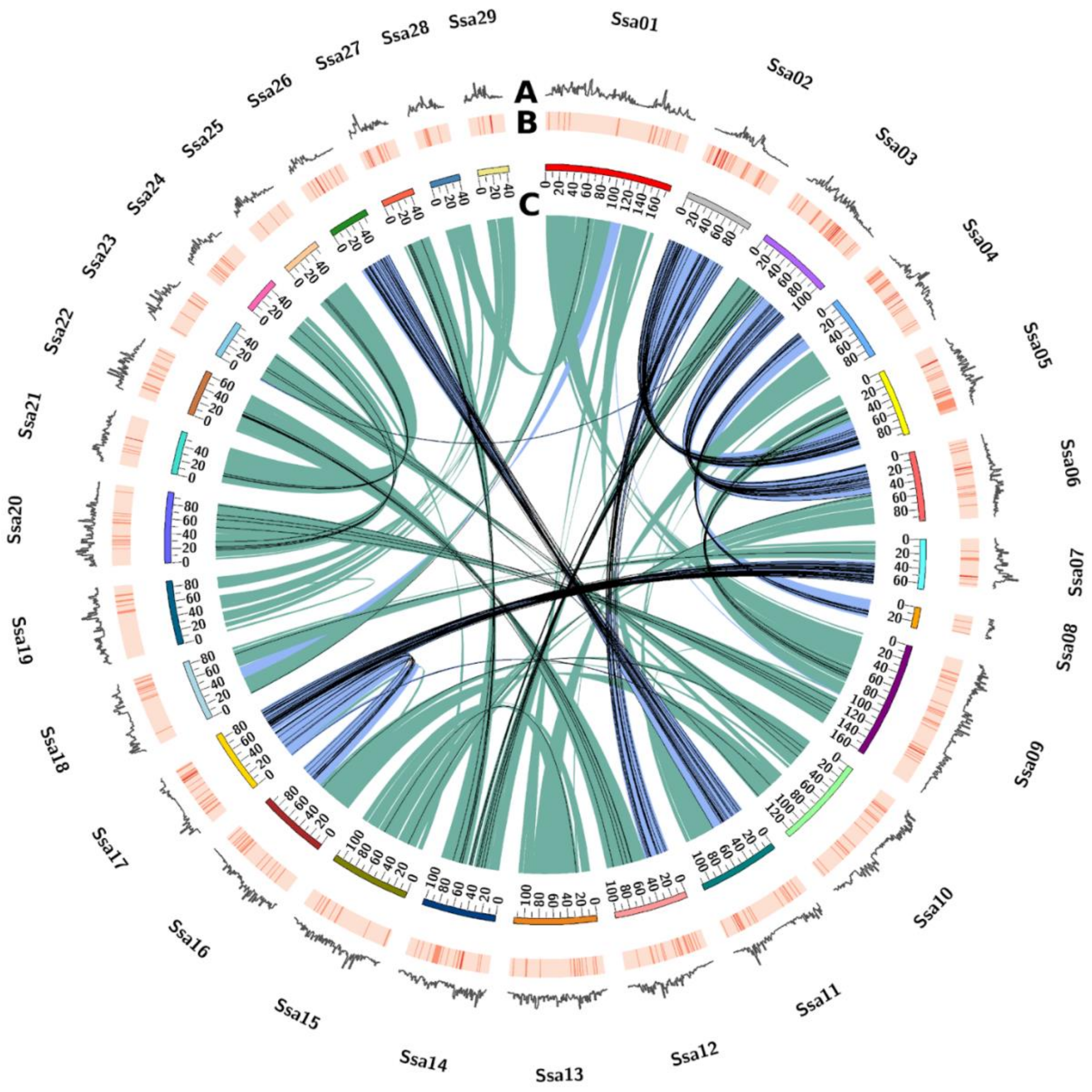


**Figure S2: Trans regulatory connections across homeologous regions**

This figure shows the distribution of trans-regulatory connections between eQTLs and genes in homeologous regions, regions that were duplicated as a result of WGD event of the Atlantic salmon genome. The outermost ring (**A**) represents the gene density across the genome, with darker areas indicating higher gene concentrations. The middle ring (**B)** shows the proportion of trans eQTLs identified within 1 Mb windows across each chromosome, where orange bars indicate eQTL density. In the inner circle (**C**), blue ribbons represent lineage-specific ohnolog resolution (LORe) regions, while green ribbons mark ancestral ohnolog resolution (AORe) regions. The black lines within the circle illustrate the trans-regulatory connections between trans-eQTLs and their associated genes, focusing on those connected across homeologous regions. Trans-regulatory connections are more frequently retained in the more recently rediploidized LORe regions compared to the older AORe regions (chi-square test, $p = 2.2e\text{-}16$)

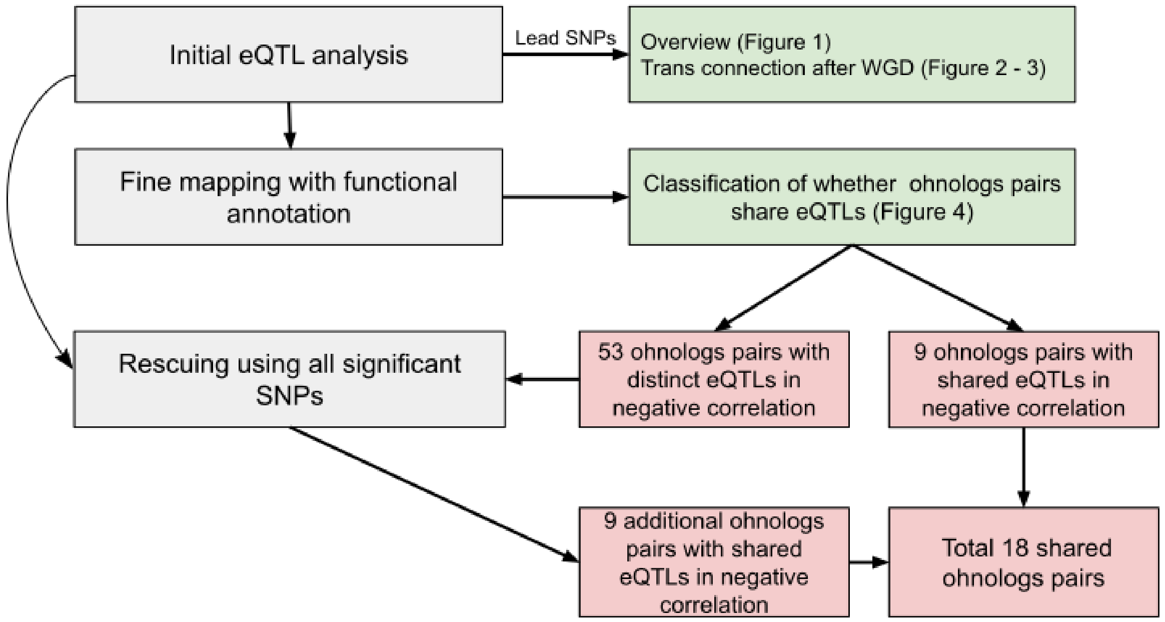


**Figure S3: Flow chart of eQTLs analysis performed**

In grey: methods used. In green: associated analysis. In red: results obtained.

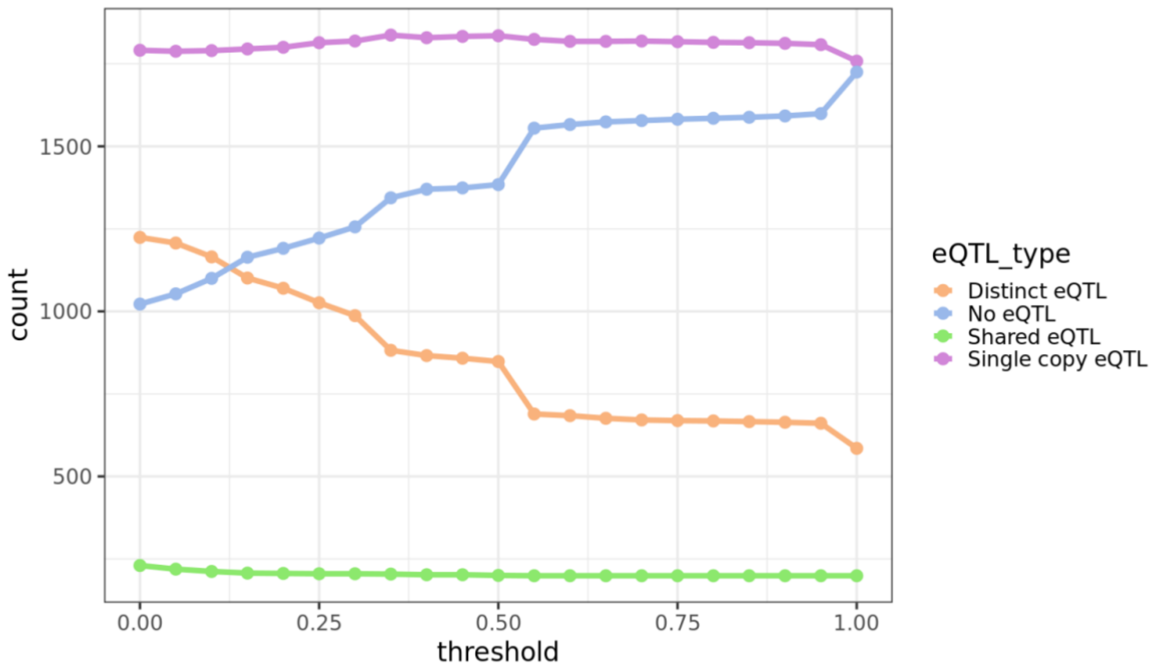


**Figure S4: Change in number of pairs in each category depending on PIP threshold**

Each dot correspond to a pip threshold tested.

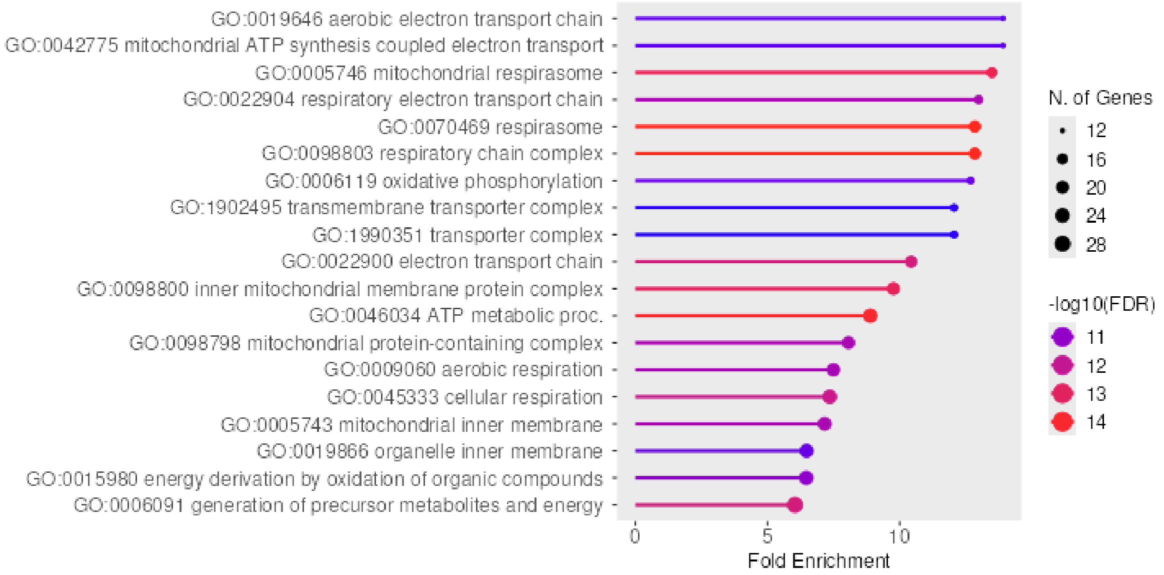


**Figure S5: GO enrichment of the 230 ohnologs with shared eQTL**

Enrichment of the 230 ohnologs using all gene set available (FDR cutoff=0.05). The 4267 ohnolog pairs were used as background, ignoring singletons.

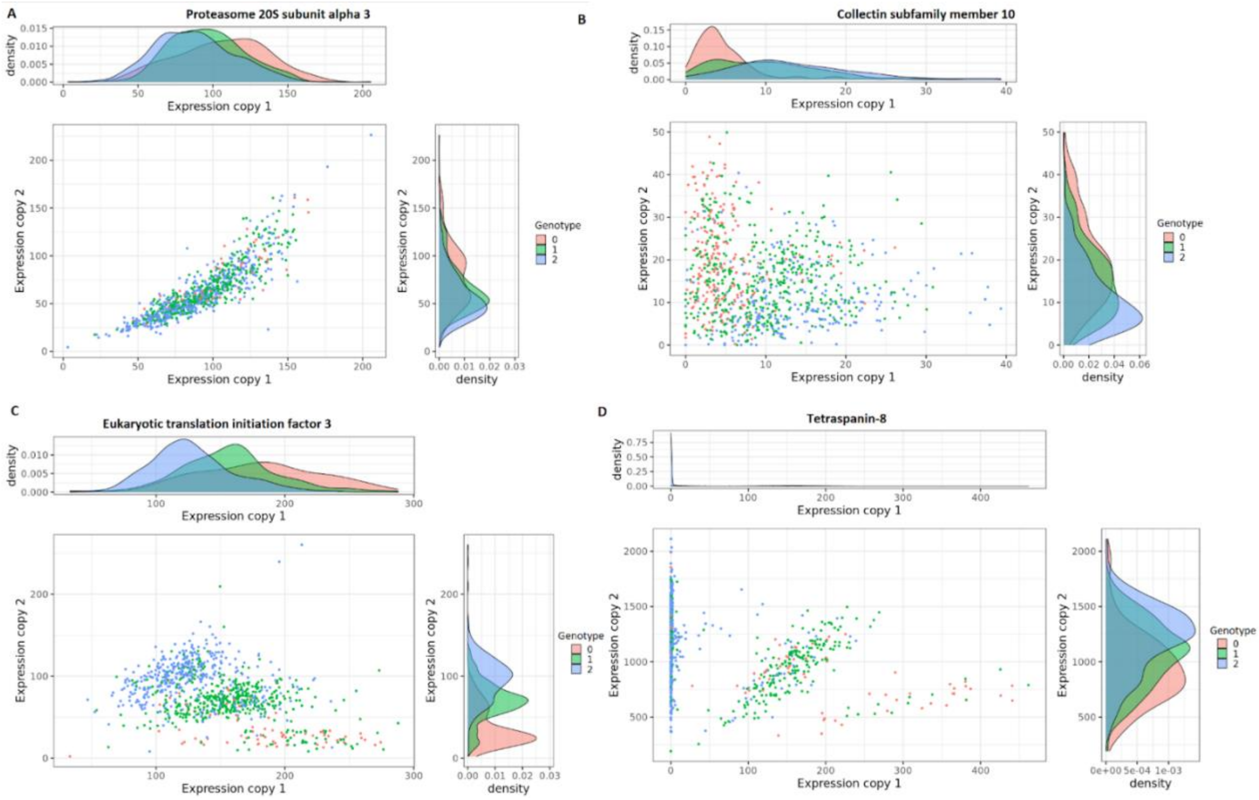


**Figure S6: Ohnologs expression according to their shared eQTL genotype**

Example of ohnologs pairs with shared eQTL, showing expression of both ohnolog copies for each individual according to their genotype for the eQTL. **A:** An example of an ohnolog pair with positive correlation, **BCD**: 3 ohnologs pairs with negative correlations. We can see 3 distinct groups of expression, representing each of the genotype

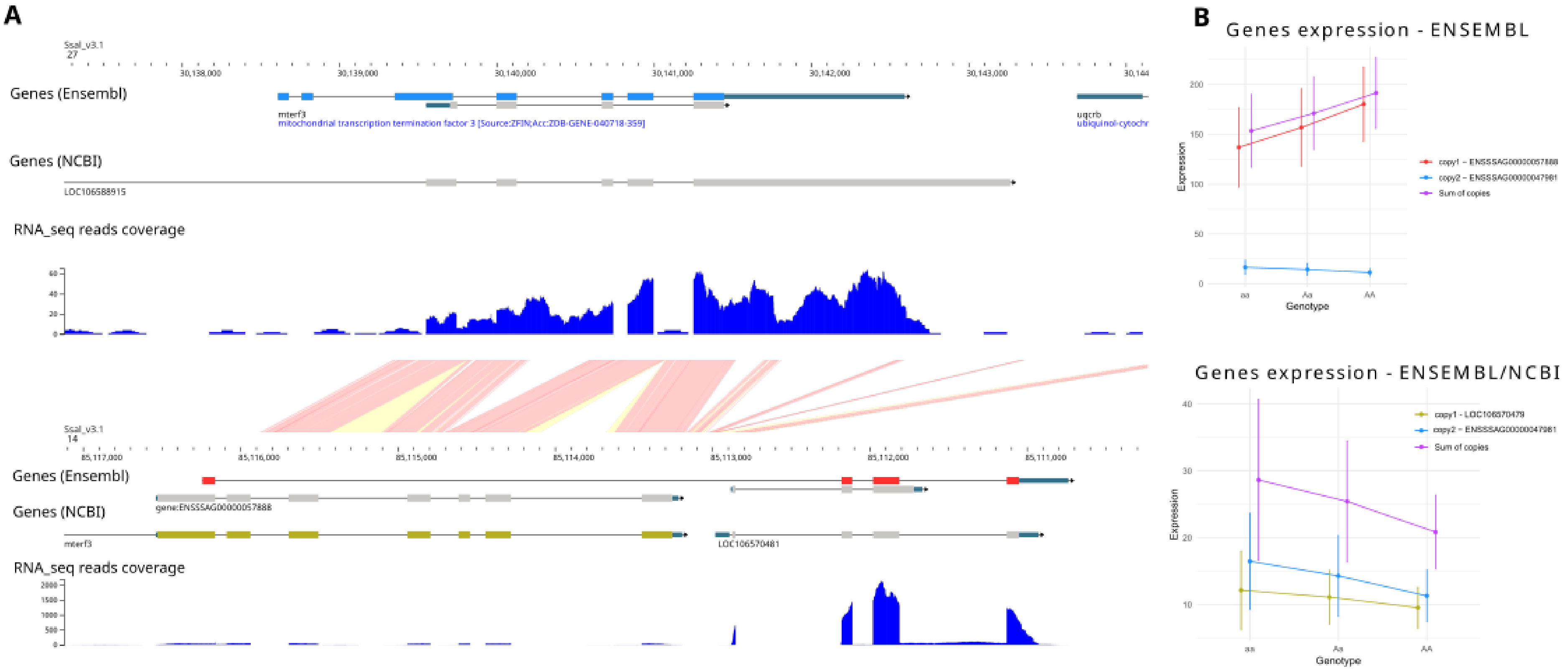


**Figure S7: Genome view and corresponding gene expression of *mterf3***

A: Genome view of chr 27 (top) and chr14 (bottom) at the location of *mterf3* ohnologs. B: Mean expression levels (in TPM) for *mterf3* gene depending on the annotation choose for *mterf3* on chr14. *Mterf3* is a gene which was found to have a shared eQTL and negative correlation between ohnologs expression. We can observe that on chr14 (bottom), annotation is discordant between NCBI and ENSEMBL. On the expression panel, we can see that the negative correlation is not observed anymore when using NCBI annotation.

**Table S1: Comparison of distribution of spearman coefficient**

Kolmogorov-smirnov test used. Threshold p-value using Bonferroni correction is 0.0083.

| | Shared eQTL | Distinct eQTL | Single copy eQTL | No eQTL |
|---|---|---|---|---|
| **Shared eQTL** | | | | |
| **Distinct eQTL** | D=0.50<br>p-val < 2.2E-16 | | | |
| **Single copy eQTL** | D=0.48<br>p-val < 2.2E-16 | D=0.03<br>p-val =0.35 | | |
| **No eQTL** | D=0.37<br>p-val < 2.2E-16 | D=0.15<br>p-val = 4.5E-12 | D=0.14<br>p-val = 1.5E-11 | |

**Table S2: Gene in dosage compensation with shared eQTL ohnologs**

P-values indicate anova test for difference in sum of expression and are corrected for multiple testing using Bonferroni correction. Significant values indicate pairs where the sum of expression is different between genotypes. When significant, the percentage of variation in expression between AA and aa genotypes is indicated in parenthesis. Expression symmetry corresponds to pairs having one copy with consistently lower expression than the others (asymmetric), or pairs with a different copy being the most expressed between each homozygous genotype (Opposite). Gene pairs that could not be classified in these categories (i.e non significantly different expression between copies in at least one genotype and no opposite pattern) are noted "NS".

| Gene | Adjusted p value | Gene 1 ID | Gene 1 location | Gene 2 ID | Gene 2 location | eQTL location | Expression Symmetry |
|---|---|---|---|---|---|---|---|
| Isocitrate Dehydrogenase (NAD(+)) 3 Non-Catalytic Subunit Beta (*idh3b*) | 5.91E-5 *** (15%) | ENSSSAG00000043068 | ssa12:50,997,002-51,006,996 | ENSSSAG00000054986 | ssa22:44,294,869-44,307,005 | ssa14:51,370,830 | Asymmetric |
| Major facilitator Superfamily Domain Containing 13A (*mfsd13al*) | 0.84 | ENSSSAG00000001778 | ssa03:77,488,542-77,494,418 | ENSSSAG00000045167 | ssa06:24,446,097-24,456,254 | ssa14:51,354,372 | NS |
| N-acyl phosphatidylethanolamine phospholipase D (*napepld*) | 0.0016 *** (12%) | ENSSSAG00000054393 | ssa07:63,480,345-63,491,382 | ENSSSAG00000117250 | ssa17:82,065,545-82,081,353 | ssa17:62,041,220 | Asymmetric |
| 2-hydroxyacyl-CoA lyase 1 (*hacl1*) | 1 | ENSSSAG00000058006 | ssa05:83,319,427-83,336,253 | ENSSSAG00000093636 | ssa02:9,545,825-9,559,847 | ssa02:14,696,160 | Opposite |
| *tetraspanin-8* | 7.70E-3 *** (9%) | ENSSSAG00000086178 | ssa17:81,256,811-81,261,845 | ENSSSAG00000087231 | ssa07:62,703,781-62,708,799 | ssa07:56,592,523 | Asymmetric |
| Carboxypeptidase M (*cpm*) | 1.60E-4 *** (18%) | ENSSSAG00000092357 | ssa17:77,867,721-77,886,108 | ENSSSAG00000109189 | ssa07:59,241,376-59,299,087 | ssa17:62,357,348 | Opposite |
| *zgc* | 0.25 | ENSSSAG00000001739 | ssa03:88,263,660-88,266,629 | ENSSSAG00000041942 | ssa06:13,783,192-13,787,468 | ssa06:12,926,921 | Asymmetric |

| | | | | | | | |
|---|---|---|---|---|---|---|---|
| Oligosaccharyltransferase complex catalytic subunit B (*stt3b*) | 1 | ENSSSAG00000002173 | ssa05:90,696,619-90,795,848 | ENSSSAG00000114008 | ssa02:1,960,032-2,083,478 | ssa05:68,517,542 | Opposite |
| Src kinase-associated phosphoprotein 2 (*skap2*) | 1 | ENSSSAG00000009565 | ssa02:9,123,961-9,143,339 | ENSSSAG00000121915 | ssa05:83,717,162-83,735,864 | ssa05:68,517,542 | Opposite |
| *collectin 10* | 1 | ENSSSAG00000011938 | ssa02:4,851,964-4,887,386 | ENSSSAG00000108018 | ssa05:87,782,557-87,817,196 | ssa05:68,654,814 | NS |
| Secretagogin (*segn*) | 1 | ENSSSAG00000036507 | ssa05:88,240,966-88,290,399 | ENSSSAG00000116949 | ssa02:4,369,425-4,417,790 | ssa05:67,379,201 | NS |
| ELOVL fatty acid elongase 8b (*elovl8a*) | 2.26E-5 *** (5%) | ENSSSAG00000044129 | ssa16:47,926,373-47,953,310 | ENSSSAG00000117896 | ssa10:12,881,358-12,886,392 | ssa14:48,488,679 | Asymmetric |
| Ribosome Binding Factor A (*rbfa*) | 1 | ENSSSAG00000083156 | ssa05:86,193,704-86,196,331 | ENSSSAG00000116300 | ssa02:6,628,362-6,630,877 | ssa02:20,664,047 | Asymmetric |
| *cc068* | 1 | ENSSSAG00000090152 | ssa02:1,387,837-1,406,608 | ENSSSAG00000110471 | ssa05:91,371,106-91,389,936 | ssa13:57,657,347 | Asymmetric |
| Protein associated with LIN7 2, MAGUK p55 family member (*PALS2*) | 0.93 | ENSSSAG00000091200 | ssa05:84,249,698-84,299,276 | ENSSSAG00000121190 | ssa02:8,502,544-8,550,092 | ssa02:55,822,073 | Asymmetric |
| Eukaryotic translation initiation factor 3, subunit M (*eif3m*) | 1 | ENSSSAG00000113858 | ssa11:38,167,566-38,174,827 | ENSSSAG00000120807 | ssa26:38,895,331-38,901,886 | ssa26:24,868,613 | Asymmetric |
